# The Temporal Relationships between White Matter Hyperintensities, Neurodegeneration, Amyloid β, and Cognition

**DOI:** 10.1101/2020.05.27.119586

**Authors:** Mahsa Dadar, Richard Camicioli, Simon Duchesne, D. Louis Collins, for the Alzheimer’s Disease Neuroimaging Initiative

**Author notes:** **Corresponding Author Information:** Mahsa Dadar, Cervo Brain Research Centre, 2601 Chemin de la Canardière, Québec, Canada, G1J 2G3. Data used in preparation of this article were obtained from the Alzheimer’s Disease Neuroimaging Initiative (ADNI) database (adni.loni.usc.edu). As such, the investigators within the ADNI contributed to the design and implementation of ADNI and/or provided data but did not participate in analysis or writing of this report. A complete listing of ADNI investigators can be found at: http://adni.loni.usc.edu/wp-ontent/uploads/how_to_apply/ADNI_Acknowledgement_List.pdf.

## Abstract

**INTRODUCTION:** Cognitive decline in Alzheimer’s disease is associated with amyloid-β accumulation, neurodegeneration and cerebral small vessel disease, but the temporal relationships between these factors is not well established.

**METHODS:** Data included white matter hyperintensity (WMH) load, grey matter (GM) atrophy and Alzheimer’s Disease Assessment Scale-Cognitive-Plus (ADAS13) scores for 720 participants and cerebrospinal fluid amyloid (Aβ1-42) for 461 participants from the Alzheimer’s Disease Neuroimaging Initiative. Linear regressions were used to assess the relationships between baseline WMH, GM, and Aβ1-42 to changes in WMH, GM, Aβ1-42, and cognition at one-year follow-up.

**RESULTS:** Baseline WMHs and Aβ1-42 predicted WMH increase and GM atrophy. Baseline WMHs, GM, and Aβ1-42 predicted worsening cognition. Only baseline Aβ1-42 predicted change in Aβ1-42.

**DISCUSSION:** Baseline WMHs lead to greater future GM atrophy and cognitive decline, suggesting that WM damage precedes neurodegeneration and cognitive decline. Baseline Aβ1-42 predicted WMH increase, suggesting a potential role of amyloid in WM damage.

**Research in Context:**

1. **Systematic Review**: Both amyloid β and neurodegeneration are primary pathologies in Alzheimer’s disease. White matter hyperintensities (indicative of presence of cerebrovascular disease) might also be part of the pathological changes in Alzheimer’s. However, the temporal relationship between white matter hyperintensities, amyloid β, neurodegeneration, and cognitive decline is still unclear.
2. **Interpretation**: Our results establish a potential temporal order between white matter hyperintensities, amyloid β, neurodegeneration, and cognitive decline, showing that white matter hyperintensities precede neurodegeneration and cognitive decline. The results provide some evidence that amyloid β deposition, in turn, precedes accumulation of white matter hyperintensities.
3. **Future Directions**: The current findings reinforce the need for future longitudinal investigations of the mechanisms through which white matter hyperintensities impact the aging population in general and Alzheimer’s disease patients, in particular.

## 1. INTRODUCTION

White matter hyperintensities^†^ (WMHs) on T2-weighted (T2w) and Fluid-attenuated inversion recovery (FLAIR) magnetic resonance images (MRIs) are indicative of the presence of cerebrovascular pathology [1]. Pathologically, WMHs have been associated with gliosis, demyelination, axonal loss, arteriosclerosis due to hypoxia, hypoperfusion, blood-brain barrier leakage, inflammation, degeneration, and amyloid angiopathy [2]. Clinically, WMHs have been associated with increased risks of cognitive decline in otherwise normal aging (NA), individuals with mild cognitive impairment (MCI), and patients with probable Alzheimer’s disease (AD) [3–8].

Accumulating evidence indicates that cerebrovascular pathology is very common in AD patients and has an important role in AD pathology, lowering the threshold for a clinical diagnosis of dementia due to AD [9]. What is less clear is whether cerebrovascular pathology occurs before, after, or at the same time as the progression of AD, and whether it has a synergistic interaction with AD pathology and neurodegeneration [9].

A number of recent studies have suggested that cerebrovascular pathology might be the starting point of a chain of events leading to AD neurodegeneration and cognitive decline [10,11]. Hypo-perfusion and ischemic changes associated with aging and vascular risk factors can result in blood supply and metabolism disturbances. Such disturbances can cause neuronal energy failure, leading to neuronal injury and acceleration in over-production and reduction in clearance of amyloid beta (Aβ), resulting in progressive cognitive deficits and neurodegeneration characteristic of AD [12,13].

On the other hand, Aβ deposition could also increase WMH burden by accelerating processes that are not necessarily vascular in nature, including neuroinflammation, reactive oxygen species production, and oxidative stress [14–16]. In this scenario, an initial rise in Aβ would damage the white matter (WM), which in turn would further elevate Aβ levels, leading to more WM damage in a cyclical process. This initial rise in Aβ would be noticeable in cerebro-spinal fluid (CSF) assays [17].

Another contributing link belongs to risk factors that are associated with WMHs and cognitive decline including hypertension, high systolic and diastolic blood pressure, hypercholesterolaemia, diabetes, obesity, high glucose levels, and smoking [2,9]. These risk factors are particularly important since they are Regarding neurodegeneration, a well-established marker of disease progression in AD is atrophy of cortical and subcortical grey matter (GM) structures. This atrophy is associated with cognitive deficits and decline in aging, MCI, and AD populations [25–29]. Whole brain measures of atrophy are strongly associated with cognitive decline and increased risk of dementia in aging, MCI, and AD [30–32].

Although it is known that both WMHs and GM atrophy contribute to cognitive deficits on the spectrum from aging to probable AD, it remains unclear whether they have an independent, synergistic, or sequential impact on cognition. In this study, we take advantage of the longitudinal data from the Alzheimer’s Disease Neuroimaging Initiative (ADNI) to investigate the temporal relationship between WMHs, GM atrophy, cognitive decline, and Aβ. Specifically, we aimed to investigate: 1) whether WMHs precede neurodegeneration and cognitive decline; and 2) whether WMHs impact Aβ progression or vice versa.

## 2. METHODS

### 2.1. Participants

We selected participants from the ADNI-1, ADNI-2, and ADNI-GO database (adni.loni.usc.edu) that had cognitive evaluations and associated T1w, T2w/PDw, and FLAIR MRIs at one-year intervals (Figure 1). The ADNI was launched in 2003 as a public-private partnership, led by Principal Investigator Michael W. Weiner, MD. The primary goal of ADNI has been to test whether serial MRI, positron emission tomography, other biological markers, and clinical and neuropsychological assessment can be combined to measure the progression of MCI and early AD. The study was approved by the institutional review board of all participating sites and written informed consent was obtained from all participants before inclusion in the study.

**Figure 1.**
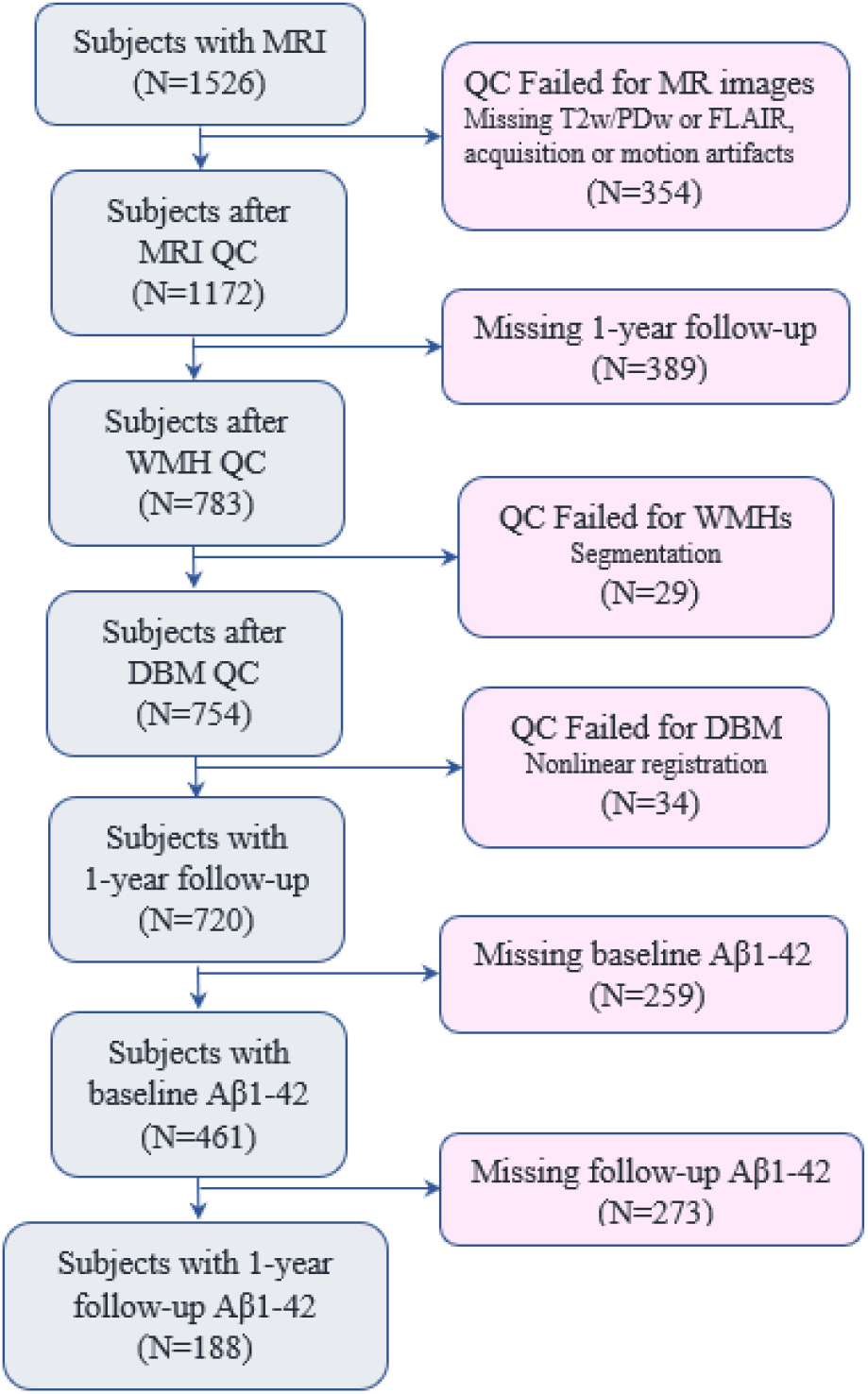
Study flowchart.

### 2.2. MRI acquisition and preprocessing

Table 1 summarizes MR imaging parameters for the data used in this study.

**Table 1 -.** Scanner information and MRI acquisition parameters for ADNI1, and ADNI2/GO datasets.

|  | Sequence | T1w | T2w/PDw | FLAIR |
| --- | --- | --- | --- | --- |
| ADNI1<br>N=421<br>N<sub>NA</sub>=124<br>N<sub>MCI</sub>=213<br>N<sub>AD</sub>=84 | Slice thickness | 1.2 mm | 3 mm |  |
|  | No. of slices | 160 | 56 |  |
|  | Field of view | 192×192 cm<sup>2</sup> | 256×256 cm<sup>2</sup> |  |
|  | Scan Matrix | 192×192 cm<sup>2</sup> | 256×256 cm<sup>2</sup> |  |
|  | Repetition time (TR) | 3000 ms | 3000/3000 ms |  |
|  | Echo time (TE) | 3.55 ms | 95.2/10.5 ms |  |
| ADNI2/GO<br>N=299<br>N<sub>NA</sub>=83<br>N<sub>MCI</sub>=183<br>N<sub>AD</sub>=33 | Slice thickness | 1.2 mm |  | 5 mm |
|  | No. of slices | 196 |  | 42 |
|  | Field of view | 256×256 cm<sup>2</sup> |  | 256×256 cm<sup>2</sup> |
|  | Scan Matrix | 256×256 cm<sup>2</sup> |  | 256×256 cm<sup>2</sup> |
|  | Repetition time (TR) | 7.2 ms |  | 11000 ms |
|  | Echo time (TE) | 3.0 ms |  | 150 ms |

T1w, T2w/PDw, and FLAIR scans were pre-processed as follows: (a) image denoising [33]; (b) intensity inhomogeneity correction; and (c) intensity scaling to a 0-100 range. For each subject, the T2w, PDw, or FLAIR scans were then co-registered to the structural T1w scan of the same timepoint using a 6-parameter rigid registration and a mutual information objective function [34]. The T1w scans were also linearly [34] and nonlinearly [35] registered to the MNI-ICBM152 unbiased average template [36].

### 2.3. WMH Measurements

Using a previously validated WMH segmentation method and a library of manual segmentations based on 100 subjects from ADNI (independent of the 720 subjects studied here), WMHs were automatically segmented at both baseline and one-year follow-up timepoints [37,38,7]. The technique uses a set of location and intensity features in combination with a random forests classifier to detect WMHs using either T1w+FLAIR or T1w+T2w/PDw images. WMH load was used as a proxy for cerebrovascular pathology and was defined as the volume of all voxels identified as WMH in the standard space (in mm^3^) and are thus normalized for head size. WMH volumes were log-transformed to achieve normal distribution.

### 2.4. GM Measurements

Deformation-based morphometry (DBM) is used to identify macroscopic anatomical changes within the population by spatially normalizing the T1w MRIs so that they all conform to the same stereotaxic space. Cross-sectional non-linear registration to the MNI-ICBM152 template resulted in a deformation field sampled on a 1mm^3^ grid for each subject/timepoint. DBM maps were calculated by taking the Jacobian determinant of the inverse deformation field [39]. Jacobian determinant reflects voxel volumes relative to the MNI-ICBM152 template; i.e. a value of 1 indicates similar volume to the same voxel in the template, and values lower/higher than one indicate volumes smaller/larger than the template. The difference in the Jacobian determinant at two timepoints can be used to estimate change in volume. A decrease in the Jacobian determinant values of a specific region between two timepoints can be interpreted as a reduced cerebral structure volume, i.e. atrophy. Using a GM mask obtained based on CerebrA atlas of the GM regions for the MNI-ICBM152 template [40], mean DBM values in the GM were calculated as whole brain measures of GM volume and used as proxies of neurodegeneration.

### 2.5. Cognitive Evaluations

All subjects received a comprehensive battery of clinical assessments and cognitive testing based on a standardized protocol (adni.loni.usc.edu) [41]. At each visit, participants underwent a series of assessments including the Alzheimer’s Disease Assessment Scale-13 (ADAS13) [42], which was used as a proxy of cognitive function.

### 2.6. Aβ Levels

CSF Amyloid β1-42 measures provided by the ADNI biomarker core (University of Pennsylvania) using microbead-based multiplex immunoassay were used to assess Aβ burden. Out of the 720 subjects, 461 and 188 had Aβ1-42 measures at baseline and follow-up visits, respectively.

### 2.7. Quality Control

Preprocessed and registered images were visually assessed for quality control (presence of imaging artifacts, failure in either linear or nonlinear registrations). WMH segmentations were also visually assessed for missing hyperintensities or over-segmentation. Either failures resulted in the participant being removed from the analyses. All MRI processing, segmentation and quality control steps were blinded to clinical outcomes. Figure 1 summarizes the QC information for the subjects that were excluded. The final sample included 720 subjects with WMH, GM volume, and ADAS13 measures available one-year intervals.

### 2.8. Statistical Analyses

Paired t-tests were used to compare baseline versus follow-up WMH, GM atrophy, ADAS13, and Aβ1-42 values. Unpaired t-tests were used to assess differences across NA, MCI, and AD groups in demographics and clinical variables. Linear regression models were used to assess the relationships between WMH load and vascular risk factors, controlling for age, sex, and WMH segmentation modality. The following linear regression models were first used to assess the relationship between baseline WMH and GM measurements, and change in WMH, GM, and cognitive function in the 720 subjects that had these measurements available:

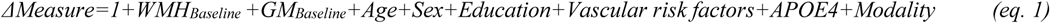

Where *ΔMeasure* indicates change in measures of interest (i.e. WMH load, GM volume, and ADAS13) between baseline and one-year follow-up visits (i.e. Measure_Follow-up_-Measure_Baseline_). *Vascular risk factors* include hypertension, systolic and diastolic blood pressure, BMI, glucose, cholesterol, and triglyceride levels. *APOE4* is a categorical variable contrasting subjects with one or two APOE4 alleles against those with zero. *Modality* is a categorical variable indicating whether WMHs were segmented using T1w+FLAIR or T1w+T2w/PDw scans to account for any potential differences in the segmentations.

A second model was used to also assess relationships with Aβ levels in the subsample that had Aβ1-42 measurements available:

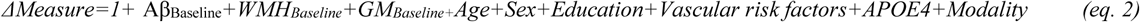

Here, *ΔMeasure* indicates change in measures of interest (i.e. WMH load, GM volume, ADAS13 and Aβ1-42). All continuous values were z-scored within the population prior to the regression analyses. The threshold for significance was *P* < 0.01, after correction for multiple comparisons using Bonferroni method.

## 3. RESULTS

### 3.1. Participants

A total of 1526 participants were available from the ADNI. After preprocessing, WMH and DBM extraction, 444 participants were removed due to failed quality control (Figure 1). The majority of these cases were missing or had significant artifacts in T2w/PD or FLAIR images. In the end, we included 720 individuals that had all MRI and clinical variables of interest available at one-year intervals (time between the two visits =1±0.1 year).

Table 2 summarizes the descriptive characteristics separately for NA, MCI, and AD participants. AD patients had significantly higher WMH load and GM atrophy levels than NA in both baseline and followup visits (*P* < 0.0003). MCI and AD groups had significantly lower CSF Aβ levels, compared to controls (*P* < 0.00001), and AD patients had significantly lower Aβ levels (*P* < 0.00001) than the MCI group. MCI and AD groups had a significantly greater proportion of subjects with APOE4 alleles (*P* < 0.001), compared to NA. There were no significant association between baseline WMHs or GM volume and APOE4 status. However, individuals with one or two APOE4 alleles had significantly higher ADAS13 scores and lower Aβ1-42 levels (*P* < 0.00001).

**Table 2.** Descriptive statistics for the participants enrolled in this study. Data are number (N) or mean ± standard deviation. *p*-values indicate group comparison results, determined by unpaired t-tests for continuous variables and χ^2^ tests for categorical variables.

| Cohort | NA | MCI | AD | MCI vs. NA | AD vs. NA | MCI vs. AD |
| --- | --- | --- | --- | --- | --- | --- |
| Number of subjects (N) | 207 | 396 | 117 | - | - | - |
| N<sub>Female</sub> | 92 | 144 | 50 | 0.07 | 0.85 | 0.25 |
| Baseline Age (years) | 75.69 ± 5.73 | 73.50 ± 7.36 | 74.49 ± 7.80 | **0.002** | 0.09 | 0.23 |
| Education (years) | 16.03 ± 2.81 | 16.11 ± 2.83 | 15.32 ± 2.82 | 0.72 | **0.03** | **0.008** |
| BMI (Kg/m<sup>2</sup>) | 26.78 ± 4.57 | 26.84 ± 4.63 | 25.94 ± 4.07 | 0.88 | 0.10 | 0.06 |
| APOE4 (0/1/2) | 142/59/6 | 192/156/48 | 37/56/24 | **0.001** | **<0.0001** | **0.002** |
| Diastolic Blood Pressure (mmHg) | 75.02 ± 10.46 | 75.38 ± 9.34 | 75.08 ± 8.86 | 0.67 | 0.96 | 0.75 |
| Systolic Blood Pressure (mmHg) | 134.96 ± 17.2 | 135.57 ± 17.1 | 135.51 ± 17.9 | 0.68 | 0.78 | 0.97 |
| Hypertension (N) | 101 | 189 | 61 | 0.87 | 0.64 | 0.46 |
| Cholesterol (mg/dL) | 193.38 ± 39.8 | 195.95 ± 40.3 | 192.18 ± 40.6 | 0.54 | 0.80 | 0.44 |
| Triglyceride (mg/dL) | 139.30 ± 89.3 | 147.17 ± 84.7 | 139.69 ± 75.3 | 0.30 | 0.97 | 0.41 |
| Glucose (mg/dL) | 99.22 ± 17.94 | 100.64 ± 23.3 | 98.12 ± 18.98 | 0.49 | 0.62 | 0.34 |
| Baseline ADAS-cog13 | 4.36 ± 9.55 | 16.82 ± 6.61 | 29.08 ± 6.49 | **<0.00001** | **<0.00001** | **<0.00001** |
| Follow-up ADAS-cog13 | 8.87 ± 4.60 | 17.69 ± 8.31 | 32.86 ± 9.34 | **<0.00001** | **<0.00001** | **<0.00001** |
| Baseline Amyloid β1-42 (pg/mL) | 198.64 ± 54.8 | 170.60 ± 51.9 | 139.31 ± 42.2 | **<0.00001** | **<0.00001** | **<0.00001** |
| Follow-up Amyloid β1-42 (pg/mL) | 202.02 ± 58.1 | 160.29 ± 50.1 | 132.13 ± 31.02 | **<0.00001** | **<0.00001** | **0.001** |
| Baseline WMH load (cm<sup>3</sup>) | 9.92 ± 13.09 | 10.15 ± 12.88 | 15.86 ± 20.83 | 0.67 | **0.0003** | **<0.00001** |
| Follow-up WMH load (cm<sup>3</sup>) | 10.79 ± 13.44 | 11.39 ± 13.90 | 18.05 ± 22.22 | 0.51 | **<0.0001** | **<0.00001** |
| Baseline GM Jacobian | 0.962 ± 0.04 | 0.958 ± 0.05 | 0.944 ± 0.04 | 0.24 | **<0.0001** | **0.005** |
| Follow-up GM Jacobian | 0.957 ± 0.04 | 0.949 ± 0.05 | 0.932 ± 0.04 | **0.03** | **<0.00001** | **0.0001** |

NA= Normal Aging. MCI= Mild Cognitive Impairment. AD= Alzheimer's Dementia. ADAS= Alzheimer's Disease Assessment Scale. WMH= White Matter Hyperintensity. CC= Cubic Centimetre. GM= Grey Matter.

### 3.2. Vascular risk factors

Controlling for age, sex, and segmentation modality, hypertension (Tstat=5.78, *P* < 0.00001), higher systolic (Tstat=2.50, *P* = 0.01) and diastolic blood pressure (Tstat=3.00, *P* = 0.002), higher glucose levels (Tstat=3.29, *P* = 0.001), and higher BMI (Tstat=2.29, *P* = 0.02) were associated with higher WMH loads at baseline. There were no differences between NA, MCI and AD groups in proportion of hypertensives, systolic blood pressure, diastolic blood pressure, BMI, glucose, cholesterol or triglyceride levels (Table 2). There was no significant association between baseline GM volume, Aβ1 −42, or ADAS13 and vascular risk factors.

### 3.3. Longitudinal comparisons

There was a significant increase in WMH load (1.23 cm^3^, 11.63% of the average baseline load) and ADAS13 scores (0.92, 5.53% of the average baseline score) indicating worsening cognitive performance, and a significant decrease in GM volume (0.0082, 0.86% of the average baseline value) indicating GM atrophy at the one-year follow-up visit compared to baseline (paired t-tests, *P* < 0.000001). There was no significant difference in Aβ1-42 levels between the baseline and follow-up visits (*P* = 0.57) for the 188 participants with available CSF results (0.97 pg/mL decrease, 0.58% of the average baseline value).

### 3.4. Relationships between baseline measurements and longitudinal change (model 1)

Baseline WMH load predicted change in WMH load over follow-up (Tstat = 6.96, *P* < 0.00001). There was no significant association between change in WMH load and age (Tstat = 0.52, *P* = 0.60), sex (Tstat = 0.78, *P* = 0.43), baseline GM volume (Tstat = −0.42, *P* = 0.67), or years of education (Tstat = −0.24, *P* = 0.80). There was no significant association with presence of APOE4 alleles or vascular risk factors. There was no significant difference across cohorts in the slopes. Figure 2 (first row) shows the relationship between age, baseline GM atrophy, and baseline WMH load and change in WMH load. Based on the model predictions, each additional 1 cm^3^ of WMHs at baseline leads to 0.71cm^3^ additional increase in WMH load during the following year, equivalent to 6.45% of the average baseline WMH load.

**Figure 2.**
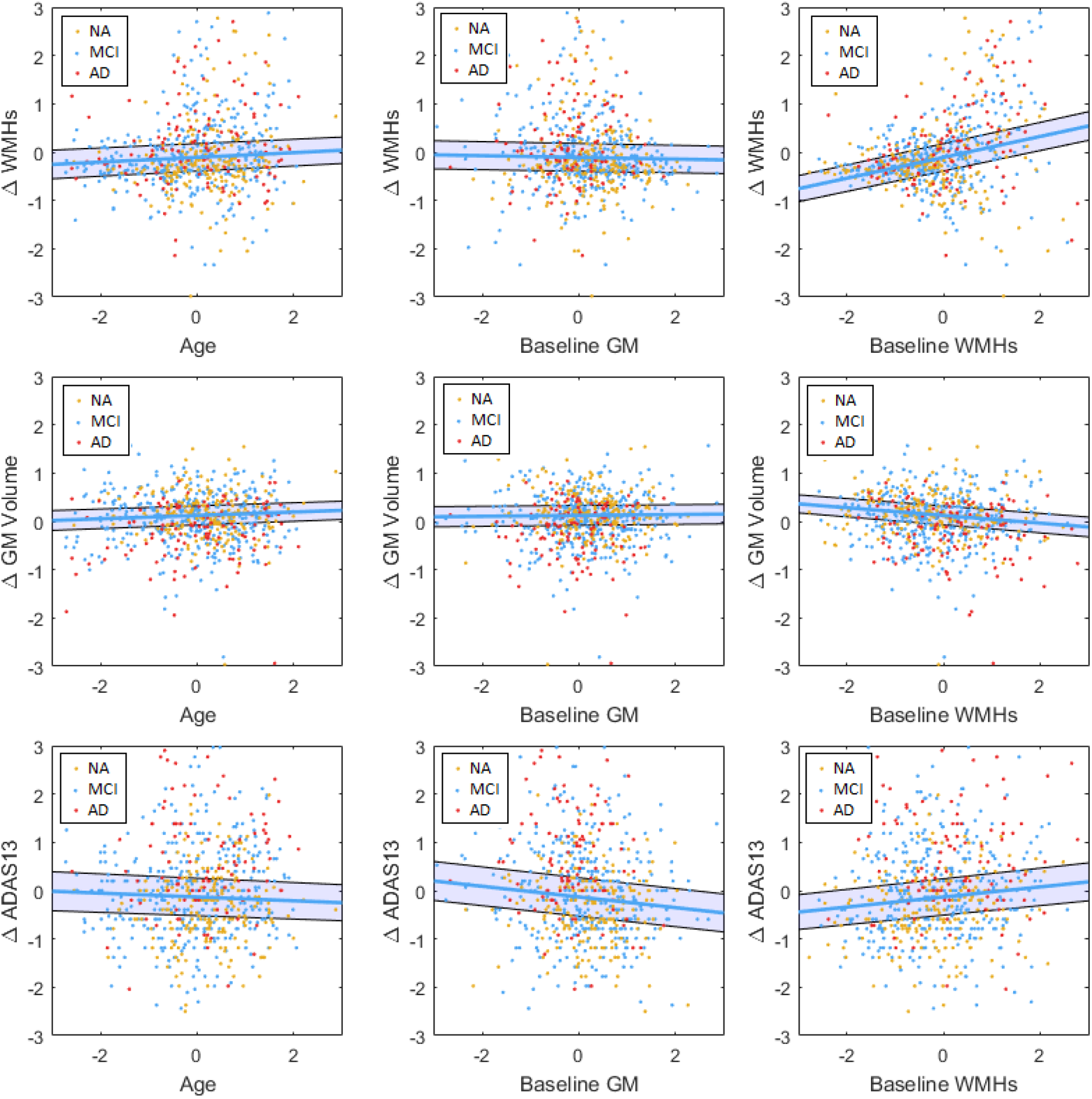
Baseline measurements and longitudinal changes in WMHs (first row), GM (second row), and ADAS13 (third row). All variables are z-scored. ΔWMH-WMH_Flollow-up_-WMH_Baseline_. ΔGM_Follow-up_-GM_Baseline_. ΔADAS13-ADAS13_Follow-up_-ADAS13_Baseline_. GM=Grey Matter. WMH= White Matter Hyperintensity. NA= Normal Aging. MCI= Mild Cognitive Impairment. AD-Alzheimer’s Dementia. ADAS= Alzheimer’s Disease Assessment Scale-13. CSF= Cerebrospinal Fluid.

Higher baseline WMH load (Tstat = −3.48, *P* = 0.0005) and presence of two APOE4 alleles (Tstat = −2.37, *P* = 0.01) predicted a decrease in GM volume during the follow-up period (i.e. atrophy). There was no significant association between age (Tstat = 1.52, *P* = 0.13), sex (Tstat = −1.05, *P* = 0.29), presence of one APOE4 allele (Tstat = −1.24, *P* = 0.21), baseline GM volume (Tstat = 0.38, *P* = 0.70), or years of education (Tstat = −0.33, *P* = 0.74) and change in GM volume. There was no significant association with vascular risk factors. There was no significant difference across cohorts in the regression slopes. Figure 2 (second row) shows the relationship between age, baseline GM volume, and baseline WMH load and change in GM volume. Based on the model predictions, each additional 1 cm^3^ of WMHs at baseline leads to 0.0014 cm^3^ additional decrease in GM during the following year, equivalent to 0.15% of the average baseline GM volume.

Both baseline WMH load and baseline GM volume predicted increases in ADAS13 scores (Tstat = 2.63, *P* = 0.008, GM Tstat = −2.42, *P* = 0.01), indicating worsening cognitive performance for individuals with higher WMH burden and lower GM volume at baseline. Presence of one (Tstat = 2.73, *P* = 0.01) or two (Tstat = 2.56, *P* = 0.01) APOE4 alleles also associated with additional worsening of cognitive performance. There was no significant association between change in ADAS13 and age (Tstat = −1.45, *P* = 0.14), sex (Tstat = 0.17, *P* = 0.76), or years of education (Tstat = −1.72, *P* = 0.09). There was no significant association with vascular risk factors. Figure 2 (third row) shows the relationship between age, baseline GM atrophy, and baseline WMH load and change in ADAS13 scores. Based on the model predictions, each additional 1 cm^3^ of WMHs at baseline leads to 0.45 additional increase in ADAS13 score during the following year, equivalent to 2.7% of the average baseline score. Similarly, a 1% lower baseline GM volume leads to an additional 0.12 increase in ADAS13 score during the following year, equivalent to 0.7% of the average baseline score; while the presence of one or two APOE4 alleles leads to an additional 1.16 and 1.70 increase in ADAS13 score during the following year, equivalent to 6.97% and 10.22% of the average baseline score, respectively.

### 3.5. Relationships with Aβ (model 2)

In the subsample of 461 participants that had Aβ1-42 data available at baseline, baseline Aβ1-42 levels were associated with change in WMH volume (Tstat = −2.07, *P* = 0.03), change in GM volume (Tstat = 1.96, *P* = 0.05), and change in ADAS13 scores (Tstat = −3.84, *P* = 0.0001). However, the relationships with WMH loads and GM volume did not survive correction for multiple comparisons. Figure 3 (first row) shows the relationship between baseline Aβ1-42 levels and change in GM, WMHs, and ADAS13, respectively. In the subsample of 188 participants that had Aβ1-42 data available at baseline and follow-up visits, change in Aβ1-42 levels was only associated with baseline Aβ1-42 (Tstat = −2.96, *P* = 0.003), and not WMH load (Tstat = −0.07, *P* = 0.94) or GM volume (Tstat = 0.11, *P* = 0.91). There was no significant association with vascular risk factors or APOE4. Figure 3 (second row) shows the relationship between change in Aβ1-42 levels and baseline Aβ1-42, GM, and WMHs, respectively.

**Figure 3.**
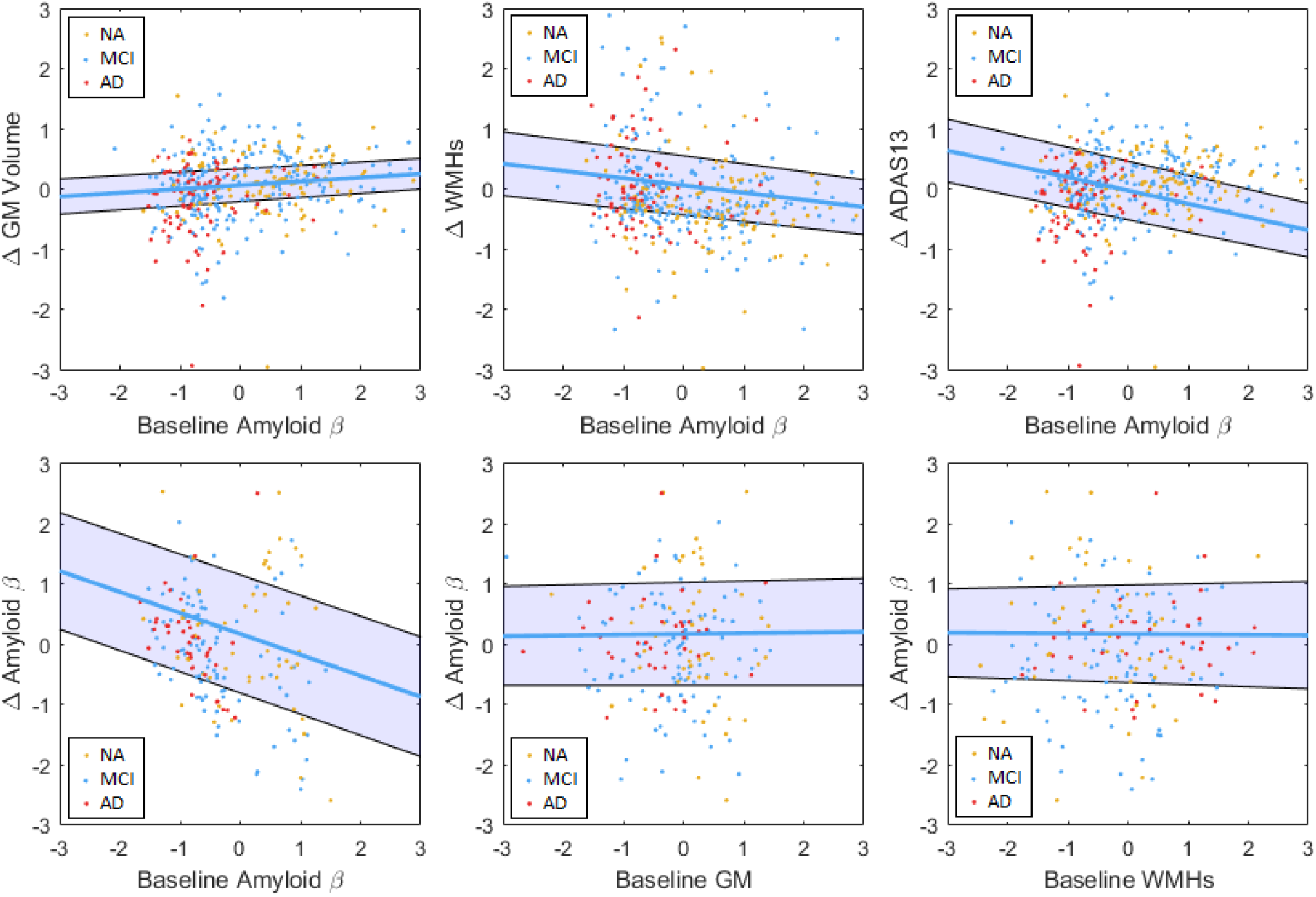
The relationship between CSF amyloid β levels and longitudinal measurements and change in CSF amyloid β levels and with baseline measurements. All values z-scored. ΔAmyloid β = Amyloid β Follow-up-Amyloid β Baseline. ΔWMH=WMH_Follow-up_-WMH_Baseline_. ΔGM=GM_Follow-up_-GM_Baseline_. ΔADAS13=ADAS13_Follow-up_-ADAS13_Baseline_. GM=Grey Matter. WMH= White Matter Hyperintensity. NA= Normal Aging. MCI= Mild Cognitive Impairment. AD= Alzheimer’s Dementia. ADAS= Alzheimer’s Disease Assessment Scale-13. CSF= Cerebrospinal Fluid.

### 3.6. Summary Model

Figure 4 summarizes the associations between APOE4, vascular risk factors, and baseline and longitudinal measurements. The arrows indicate significant associations based on the analyses in the previous sections.

**Figure 4.**
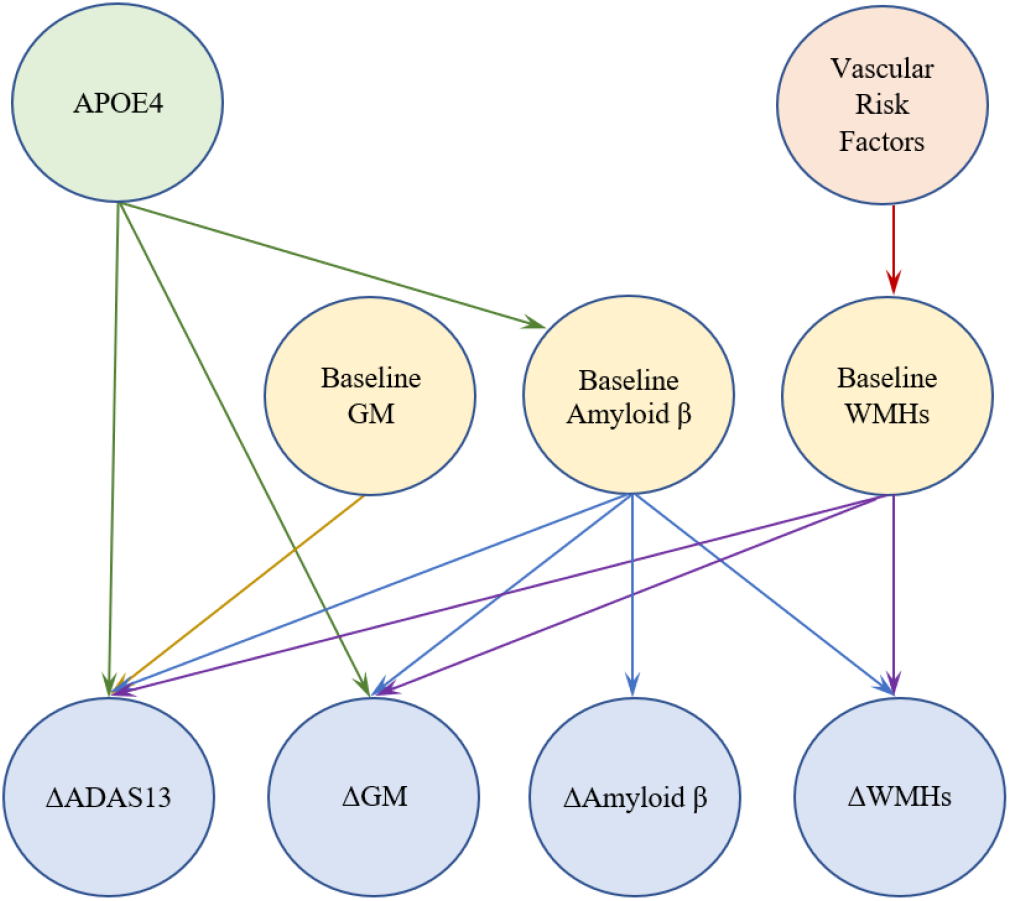
The model summarizing the relationships between APOE4, vascular risk factors, and baseline and longitudinal measurements. All values z-scored. ΔAmyloid β = Amyloid β Follow-up-Amyloid β Baseline. ΔWMH=WMHFollow-up-WMHBaseline. ΔGM=GMFollow-up-GMBaseline. ΔADAS13=ADAS13Follow-up-ADAS13Baseline. GM=Grey Matter. WMH= White Matter Hyperintensity. ADAS= Alzheimer’s Disease Assessment Scale-13.

## 4. DISCUSSION

In this study, we investigated the temporal relationships between WMHs, GM volume, Aβ, and cognitive performance in a cohort of cognitively healthy aging, MCI, and probable AD individuals. Our results showed a contribution of baseline WMH burden and Aβ to increase in WMHs, GM atrophy and cognitive decline (Figure 4).

### 4.1. Findings

There was a significant association between baseline WMH load and decrease in GM volume (i.e. atrophy), while change in WMH load was not associated with baseline GM volume (Figure 4). These results indicate that WMHs might precede GM atrophy. Taken together with the fact that WMH progression can be slowed down and possibly even prevented through anti-hypertensive mediations and lifestyle changes [43], this is an important finding, raising the possibility for intervention before irreversible neurological damage occurs. This is also in line with the studies showing that reduction of vascular disease risk and WMHs decreases the risk of cognitive deterioration [18–24].

Change in ADAS13 scores (indicating worsening cognitive performance) was significantly associated with baseline WMH load, GM volume, Aβ levels, and presence of one or two APOE alleles (Figure 4). These findings are also in line with previous studies reporting an impact of WMH burden [3–6,8,7] and GM atrophy on cognitive performance/decline [31,32]. However, those studies had not assessed the impact of WMHs and GM atrophy simultaneously. Billelo et al. reported a contribution of both WMHs and GM atrophy in preselected regions of interest to cognitive decline measured by the Consortium to Establish a Registry for AD (CERAD) scores in a similar but smaller (N=158) sample, however, they did not assess WMH and GM atrophy measures in the same model [44]. Similarly, in a cohort of 65 non-demented elderly individuals, van der Flier et al. reported an association between both WMH burden and whole brain atrophy and decline in the Cambridge Cognitive Examination (CAMCOG) scores [30]. In this study, using a much larger sample (N=720) and controlling for vascular risk factors and baseline Aβ levels (in a subset of 461 individuals), we were able to show a contribution of both GM and WM pathology to worsening cognitive performance.

Finally, controlling for vascular risk factors, we observed an association trend between lower baseline CSF Aβ1 −42 levels and increase in WMH loads, lending support to the hypothesis that Aβ deposition in the brain could increase WMH burden by accelerating processes that are not necessarily vascular in nature, such as neuroinflammation and oxidative stress [14–16,45]. On the other hand, baseline WMH loads were not associated with change in Aβ1-42 levels. However, since the sample including follow-up Aβ1-42 values was significantly smaller (188 versus 461), this finding should be interpreted with caution. In addition, the participants that had Aβ1-42 testing might have different characteristics than those that did not. In fact, a marginally higher proportion of the AD subjects than NA (*P* = 0.07) and MCI (*P* = 0.06) had Aβ1-42 values available, leading to significantly lower GM volumes in this subsample (*P* = 0.0003). There was no significant difference in WMHs or ADAS13 scores.

### 4.2. Strengths and limitations

The image processing, registration, and segmentation methods used were all developed and extensively validated in multi-center/scanner datasets, and have since been used in many such studies [46,8,34,47,38,48–50].

We used a relatively short follow-up term (one-year) to assess change in MRI and clinical measures. Although this might prevent us from detecting more extensive levels of change, it allowed us to capture the more subtle changes that occur in a shorter duration. Although a longer follow-up would likely allow us to observe greater associations between the variables, it might also obfuscate the temporal relationships due to the prolonged co-existence of the pathologies. In addition, we were able to include a larger number of subjects with all the MRI and clinical measurements available for the follow-up period.

WMHs were segmented using T1w+FLAIR and T1w+T2w/PDw scans in ADNI1 and ADNI2/GO data, respectively. To ensure that differences in FLAIR versus T2w/PDw characteristics did not affect the results, we performed an experiment segmenting WMHs in 70 cases that had T2w/PDw and FLAIR scans, using either T1w+FLAIR or T1w+T2w/PDw scans, respectively. The obtained volumes had a very high correlation (r=0.97, *P* < 0.00001), and were not significantly different (*P* = 0.65, paired t-test). In addition, we included segmentation modality as a covariate to account for any remaining differences.

The ADNI database is a cohort of relatively well-educated individuals with good access to medical care. While this relative homogeneity allows for investigation of MRI and clinical changes without excess confounds, it might not be representative of other populations with lower socioeconomic status where access to health services are more limited and a higher degree of vascular risk factors are generally present. Further investigations in more representative cohorts are necessary to observe the full spectrum of associations.

## 5. CONCLUSION

Understanding the temporal relationships between WMHs, GM atrophy, Aβ, and cognitive decline might elucidate some of the underlying mechanisms of cognitive decline in the aging population. Our results suggest that a higher WMH load at baseline might lead to greater future GM atrophy and decline in cognitive performance, indicating that earlier WM damage might precede neurodegeneration and cognitive decline. Furthermore, we observed an impact of baseline Aβ levels on increase in WMH loads, independent of vascular risk factors, indicating that (a portion of) the WMHs observed in AD patients might result from Aβ deposition and AD related pathologies.

## 7. ACKNOWLEDGEMENTS

Data collection and sharing for this project was funded by the Alzheimer’s Disease Neuroimaging Initiative (ADNI) (National Institutes of Health Grant U01 AG024904) and DOD ADNI (Department of Defense award number W81XWH-12-2-0012). ADNI is funded by the National Institute on Aging, the National Institute of Biomedical Imaging and Bioengineering, and through generous contributions from the following: AbbVie, Alzheimer’s Association; Alzheimer’s Drug Discovery Foundation; Araclon Biotech; BioClinica, Inc.; Biogen; Bristol-Myers Squibb Company; CereSpir, Inc.; Cogstate; Eisai Inc.; Elan Pharmaceuticals, Inc.; Eli Lilly and Company; EuroImmun; F. Hoffmann-La Roche Ltd and its affiliated company Genentech, Inc.; Fujirebio; GE Healthcare; IXICO Ltd.; Janssen Alzheimer Immunotherapy Research & Development, LLC.; Johnson & Johnson Pharmaceutical Research & Development LLC.; Lumosity; Lundbeck; Merck & Co., Inc.; Meso Scale Diagnostics, LLC.; NeuroRx Research; Neurotrack Technologies; Novartis Pharmaceuticals Corporation; Pfizer Inc.; Piramal Imaging; Servier; Takeda Pharmaceutical Company; and Transition Therapeutics. The Canadian Institutes of Health Research is providing funds to support ADNI clinical sites in Canada. Private sector contributions are facilitated by the Foundation for the National Institutes of Health (www.fnih.org). The grantee organization is the Northern California Institute for Research and Education, and the study is coordinated by the Alzheimer’s Therapeutic Research Institute at the University of Southern California. ADNI data are disseminated by the Laboratory for Neuro Imaging at the University of Southern California.

## Footnotes

† Abbreviations: Aβ: Amyloid beta; AD: Alzheimer’s disease; ADAS13: Alzheimer’s Disease Assessment Scale-13; CSF: Cerebrospinal fluid; DBM: Deformation-based morphometry; FLAIR: Fluid attenuated inversion recovery; GM: Grey matter; MCI; Mild cognitive impairment; MRI: Magnetic resonance imaging; NA: normal aging; T2w: T2-weighted; WM: White matter; WMH: White matter hyperintensities amenable to prevention and treatment [9] and their reduction might impede WMH progression and cognitive deterioration [18–24].

